# Ether lipid and sphingolipid expression patterns are G-protein coupled estrogen receptor 1-dependently altered in breast cancer cells

**DOI:** 10.1101/2020.07.20.212894

**Authors:** Lisa Hahnefeld, Lisa Gruber, Nina Schömel, Caroline Fischer, Peter Mattjus, Robert Gurke, Martina Beretta, Nerea Ferreirós, Gerd Geisslinger, Marthe-Susanna Wegner

**Affiliations:** *pharmazentrum frankfurt*/ZAFES, Institute of Clinical Pharmacology, Johann Wolfgang Goethe University, Theodor Stern-Kai 7, 60590 Frankfurt am Main, Germany; Åbo Akademi University, Biochemistry, Faculty of Science and Engineering Artillerigatan 6A, III, BioCity Fl-20520 Turku, Finland; Fraunhofer Institute for Molecular Biology and Applied Ecology IME, Branch for Translational Medicine and Pharmacology TMP, Theodor Stern-Kai 7, 60590 Frankfurt am Main, Germany; School of Biotechnology and Biomolecular Sciences, University of New South Wales, Sydney, New South Wales 2052, Australia

**Keywords:** sphingolipids, ether lipids, GPER1, AGMO, AGPS

## Abstract

Identifying co-expression of lipid species is challenging, but indispensable to identify novel therapeutic targets for breast cancer treatment. Lipid metabolism is often dysregulated in cancer cells, and changes in lipid metabolism affect cellular processes such as proliferation, autophagy, and tumor development. In addition to mRNA analysis of sphingolipid metabolizing enzymes, we performed liquid chromatography time-of-flight mass spectrometry analysis in three breast cancer cell lines. These breast cancer cell lines differ in estrogen receptor and G-protein coupled estrogen receptor 1 status. Our data show that sphingolipids and non-sphingolipids are strongly increased in SKBr3 cells. SKBr3 cells are estrogen receptor negative and G-protein coupled estrogen receptor 1 positive. Treatment with G15, a G-protein coupled estrogen receptor 1 antagonist, abolishes the effect of increased sphingolipid and non-sphingolipid levels in SKBr3 cells. In particular, ether lipids are expressed at much higher levels in cancer compared to normal cells and are strongly increased in SKBr3 cells. Our analysis reveals that this is accompanied by increased sphingolipid levels such as ceramide, sphingadiene-ceramide and sphingomyelin. This shows the importance of focusing on more than one lipid class when investigating molecular mechanisms in breast cancer cells. Our analysis allows unbiased screening for different lipid classes leading to identification of co-expression patterns of lipids in the context of breast cancer. Co-expression of different lipid classes could influence tumorigenic potential of breast cancer cells. Identification of co-regulated lipid species is important to achieve improved breast cancer treatment outcome.

**Highlights:**

- LC-HRMS analysis allows identification of co-expression between lipid classes
- Putative co-expression of sphingolipid and non-sphingolipid classes
- Ether lipids are strongly upregulated in SKBr3 cells (ER negative, GPER1 positive)

## 1. Introduction

Breast cancer is the most common cancer among females in North America, Europe and Oceania and shares the lead with cervical cancers in South America, Africa and most of Asia (reviewed in (Torre et al., 2016)). 70 to 78 % of the breast tumors express the transcriptionally more active *estrogen receptor* (ER) subtype α, which is declared as an ER positive (+) status (Pujol et al., 1994, Chu and Anderson, 2002). ER + tumor proliferation is hormone driven. Current therapies are based on either lowering patient estrogen levels by aromatase inhibition or by blocking estrogen-mediated signaling pathways through *selective ER modulator* (SERM) or *selective ER downregulator* (SERD) treatment. Normally, following diffusion into the cell, estrogen mediates its function by binding on ERs, which leads to translocation of the receptor into the nucleus. This results in activation of manifold signaling pathways by gene transcription alteration. Membrane-associated ERs also include *G-protein coupled estrogen receptor 1* (GPER1). GPER1 mediates rapid non-genomic as well as indirect genomic responses (reviewed in (Hsu et al., 2019)). This membrane-associated ER is involved in physiological processes such as cell growth and pathophysiological processes such as tumor development (reviewed in (Olde and Leeb-Lundberg, 2009, Wang et al., 2010)) and is discussed controversially in the literature. For example, it is still unclear where exactly GPER1 is subcellularly located and whether this receptor contributes to tumorigenic potential, or indicates less aggressiveness of breast cancer cells.

It has been shown that abnormalities of lipids influence cellular processes resulting in metabolic disorders or tumor development (reviewed in (Long et al., 2018, Pakiet et al., 2019)). Thereby, the influence of lipids ranges from promoting cancer to suppressing cancer (reviewed in (Lim, 2018)). It has been observed that ether lipids are expressed at much higher levels in cancer as compared to normal cells (reviewed in (Dean and Lodhi, 2018)). Also, plasma ether-linked *phosphocholine* (PC) species, which are elevated in breast cancer patients, can be used as a biomarker for the diagnosis of breast cancer (Chen et al., 2016). Furthermore, Benjamin et al. showed elevated ether lipid levels in more aggressive breast cancer cell lines such as 231 MFP (Benjamin et al., 2013). It has also been speculated that ether lipids seem to be capable of taking over the role of certain sphingolipids both in cells and organisms (reviewed in (Jimenez-Rojo and Riezman, 2019)). Furthermore, deregulated sphingolipid metabolism contributes to tumor development and progression (reviewed in (Ryland et al., 2011, Wegner et al., 2016)). Sphingolipids such as ceramide and sphingomyelin are significantly increased in human breast cancer (Nagahashi et al., 2016) compared to normal tissue (reviewed in (Furuya et al., 2011)). However, there is contradictory data relating to the involvement of lipid classes in cellular pathophysiological processes. One example is the finding that inhibition or silencing of *acetyl-CoA carboxylase* (ACC), which is essential for fatty acid synthesis, is shown to limit cancer cell growth (reviewed in (Lim, 2018)). Surprisingly, liver-specific ACC knockout leads to increased tumor incidence in a hepatocellular carcinogen *diethylnitrosamine* (DEN) mouse model (Nelson et al., 2017). This indicates that lipogenesis is not essential for liver tumorigenesis in the DEN mouse model. Another example is *adipose triglyceride lipase* (ATGL), which inhibits growth of several cancer cell types when suppressed, but mice lacking ATGL develop lung tumors (reviewed in (Chen and Huang, 2019)). Given the complexity of the influence that lipids have on tumor development, it is important to analyze biological samples by a method which allows an unbiased investigation of several species of lipids instead of focusing on a single lipid class.

Here, we investigated co-expression of different lipid species in breast cancer cells with differing ER and GPER1 status using *liquid chromatography time-of-flight mass spectrometry* (LC-HRMS). Our LC-HRMS analysis show co-regulation of sphingolipid and non-sphingolipid expression and indicate that tumorigenic potential of breast cancer cells is affected by this. Strikingly, ether lipids and sphingolipids are strongly increased in SKBr3 cells (ER negative, GPER1 positive). This GPER1-dependent co-regulation of sphingolipid and non-sphingolipid expression is a novel finding, which might contribute to the identification of novel therapeutic targets in breast cancer therapy.

## 2. Results

### 2.1. *Estrogen receptor* (ER) and *G-protein coupled estrogen receptor 1* (GPER1) status of breast cancer cells

First we analyzed *estrogen receptor* (ER) and *G-protein coupled estrogen receptor* 1 (GPER1) mRNA expression status of T47D, MCF-7 and SKBr3 cells by *quantitative RealTime* (qRT)-PCR. T47D cells only express ERα resulting in an ER + and GPER1 – status **(Figure 1A).** MCF-7 cells are ERα, ERβ and GPER1 expressing breast cancer cells. Accordingly, MCF-7 cells are identified to exhibit an ER + and GPER1 + status. GPER1 mRNA expression is the highest in SKBr3 cells as compared to the other cell lines showing a GPER1 + status, whereas ERα and ERß are rarely detectable. Therefore, SKBr3 cells are an ER - status breast cancer cell line. Relative mRNA expression below the value of 200 is assumed to be a negative status for the respective gene. The results displayed in **Figure 1A** are in line with other studies (Mota et al., 2017, Deng et al., 2020).

**Figure 1:**
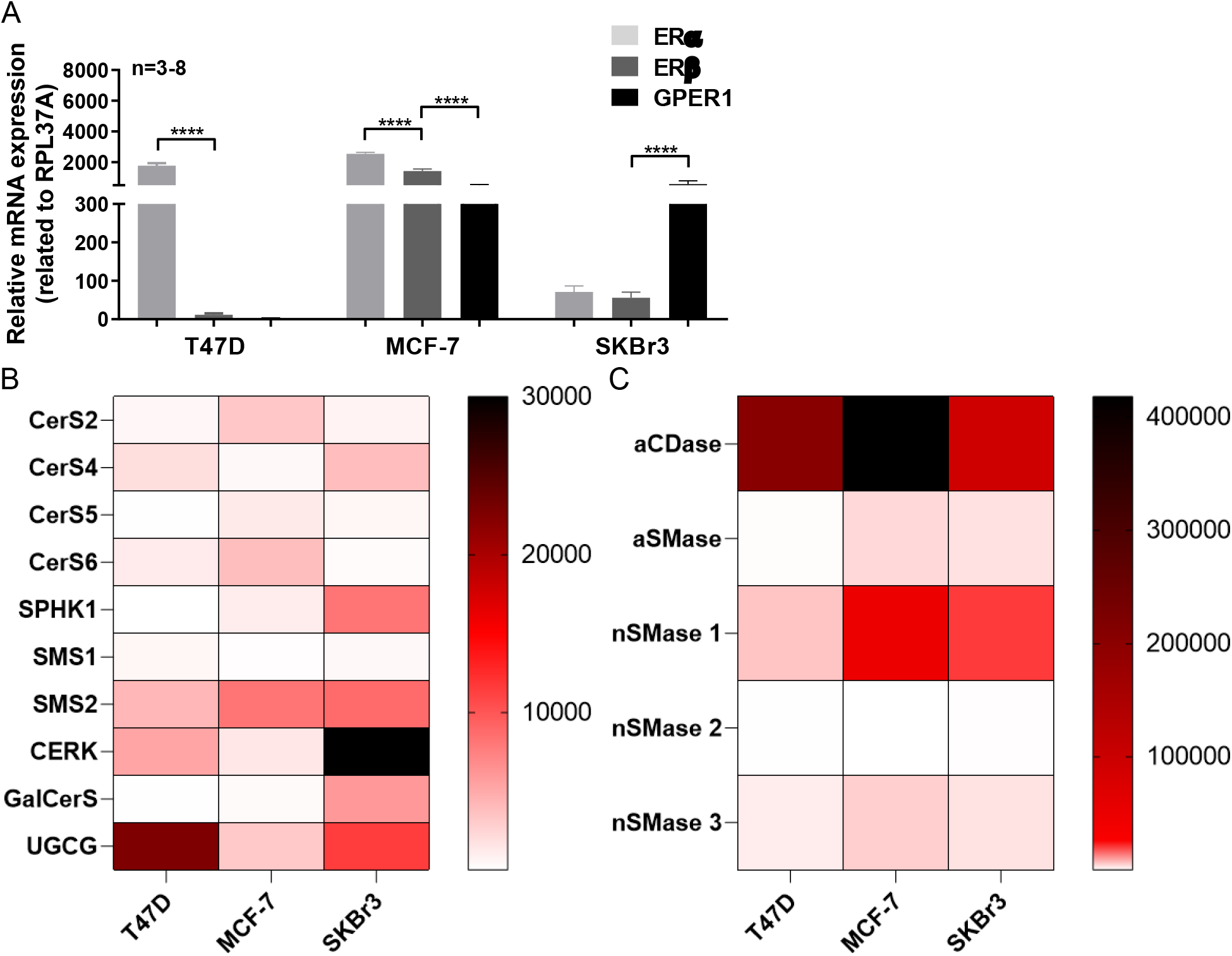
*Estrogen receptor* (ER) and *G-protein coupled estrogen receptor 1* (GPER1) status and mRNA expression analysis of sphingolipid metabolizing enzymes in breast cancer cells by qRT-PCR. **A** ERα, ERβ and GPER1 mRNA expression related to the housekeeping gene RPL37A. Data are presented as a mean of n=3-8 ± *standard error of the mean* (SEM). Tukey’s multiple comparison test. \*\*\*\**p≤0.0001*. **B** Heatmap of anabolic sphingolipid metabolizing enzyme mRNA expression. mRNA expression is related to the housekeeping gene RPL37A. High values are represented as *black*, middle range values are *red* and low values are shown in *white*. Data are presented as a mean of *n=3. Ceramide synthase X* (CerS X), *sphingosine kinase 1* (SPHK1), *sphingomyelin synthase 1 and 2* (SMS1 and 2), *ceramide kinase 1* (CERK), *galactosylceramide synthase* (GalCerS), *UDP-glucose ceramide glucosyltransferase* (UGCG). **C** Heatmap of catabolic sphingolipid metabolizing enzyme mRNA expression. mRNA expression is related to the housekeeping gene RPL37A. High values are represented as *black*, middle range values are *red* and low values are shown in *white*. Data are presented as a mean of n=3. *Acid ceramidase* (aCDase), *acid sphingomyelinase* (aSMase), *neutral sphingomyelinase 1, 2* and *3* (nSMase 1, 2 and 3).

### 2.2. Basal mRNA expression of sphingolipid metabolizing enzymes

Next we determined the basal mRNA expression of several anabolic and catabolic sphingolipid metabolizing enzymes in T47D, MCF-7 and SKBr3 cells. In general, *sphingomyelin synthase 2* (SMS2), *ceramide kinase* (CERK), *UDP-glucose ceramide glucosyltransferase* (UGCG), *acid ceramidase* (aCDase) and *neutral sphingomyelinase 1* (nSMase 1) are the most highly expressed genes in all three breast cancer cell lines **(Figure 1B and C**). Exceptions are SPHK1 and GalCerS, which are highly expressed in SKBr3, but not T47D and MCF-7 cells **(Figure 1B).** Compared to MCF-7 and SKBr3 cells, T47D cells express higher amounts of UGCG **(Figure 1B).** MCF-7 cells as compared to the other cell lines exhibit a strong mRNA expression of the anabolic enzymes *ceramide synthase 2* and *6* (CerS2 and CerS6) **(Figure 1B).** Notably, UGCG mRNA expression is comparable to CerS2 and CerS6 mRNA expression in MCF-7 cells and SMS2 is the most highly expressed gene in MCF-7 cells **(Figure 1B).** The catabolic enzymes aCDase and nSMase 1 are also highly expressed in MCF-7 cells when compared to the other cell lines **(Figure 1C).** Since MCF-7 cells express ER subtypes α and β and GPER1 these enzymes could potentially be regulated by ERα, β and GPER1. Interestingly, *ceramide synthase 4* (CerS4), *sphingosine kinase 1* (SPHK1), CERK and *galactosylceramide synthase* (GalCerS) are strongly expressed in GPER1 + SKBr3 cells compared to the other cell lines **(Figure 1B).** The data reveal differing expression patterns of sphingolipid metabolizing enzymes in breast cancer cells exhibiting unequal ER and GPER1 status.

### 2.3. ER-and GPER1-dependent sphingolipid levels

In order to screen for ER-dependent changes on sphingolipid levels we performed LC-HRMS analysis. ER - SKBr3 cells exhibit significantly reduced *dihydroceramide* (dhCer) levels compared to MCF-7 cells **(Figure 2A** and **supplemental data 1A),** whereas ceramide levels are strongly increased **(Figure 2B** and **supplemental data 1A).** Sphingadiene-ceramide concentration is also strongly increased in SKBr3 cells compared to ER + cells, albeit MCF-7 cells exhibit a higher sphingadiene-ceramide concentration than T47D cells **(Figure 2C** and **supplemental data 1A).** *Galactosylceramide* (GalCer)/*glucosylceramide* (GlcCer) levels in SKBr3 cells are increased compared to MCF-7 cells, whereas no significant differences between MCF-7 and T47D cells were detected **(Figure 2D** and **supplemental data 1B).** *Lactosylceramide* (LacCer) concentration in T47D cells is the lowest compared to MCF-7 and SKBr3 cells **(Figure 2E** and **supplemental data 1B).** Additionally, *sphingomyelin* (SM) concentration appears to follow the expression of GPER1-mRNA with highest level in SKBr3 cells and significantly lowest level in T47D cells **(Figure 2F** and **supplemental data 1C).** Treatment with G15, a GPER1 antagonist, significantly reduced ceramide and sphingadiene-ceramide level in SKBr3 cells. Dihydroceramide, ceramide, GalCer/GlcCer, LacCer and sphingomyelin levels are significantly increased in MCF-7 cells **(supplemental data 1D).** No effect of G15 treatment on the analyzed lipid levels in T47D (GPER1 -) could be detected **(supplemental data 1D** and **2H).** In summary, sphingolipid analysis revealed that the high anabolic enzyme mRNA expression in SKBr3 cells is reflected on sphingolipid levels by high ceramide, sphingadiene-ceramide, GalCer/GlcCer and SM levels. Furthermore, the G15 treatment data indicate that sphingolipid levels are GPER1-dependently regulated.

**Figure 2:**
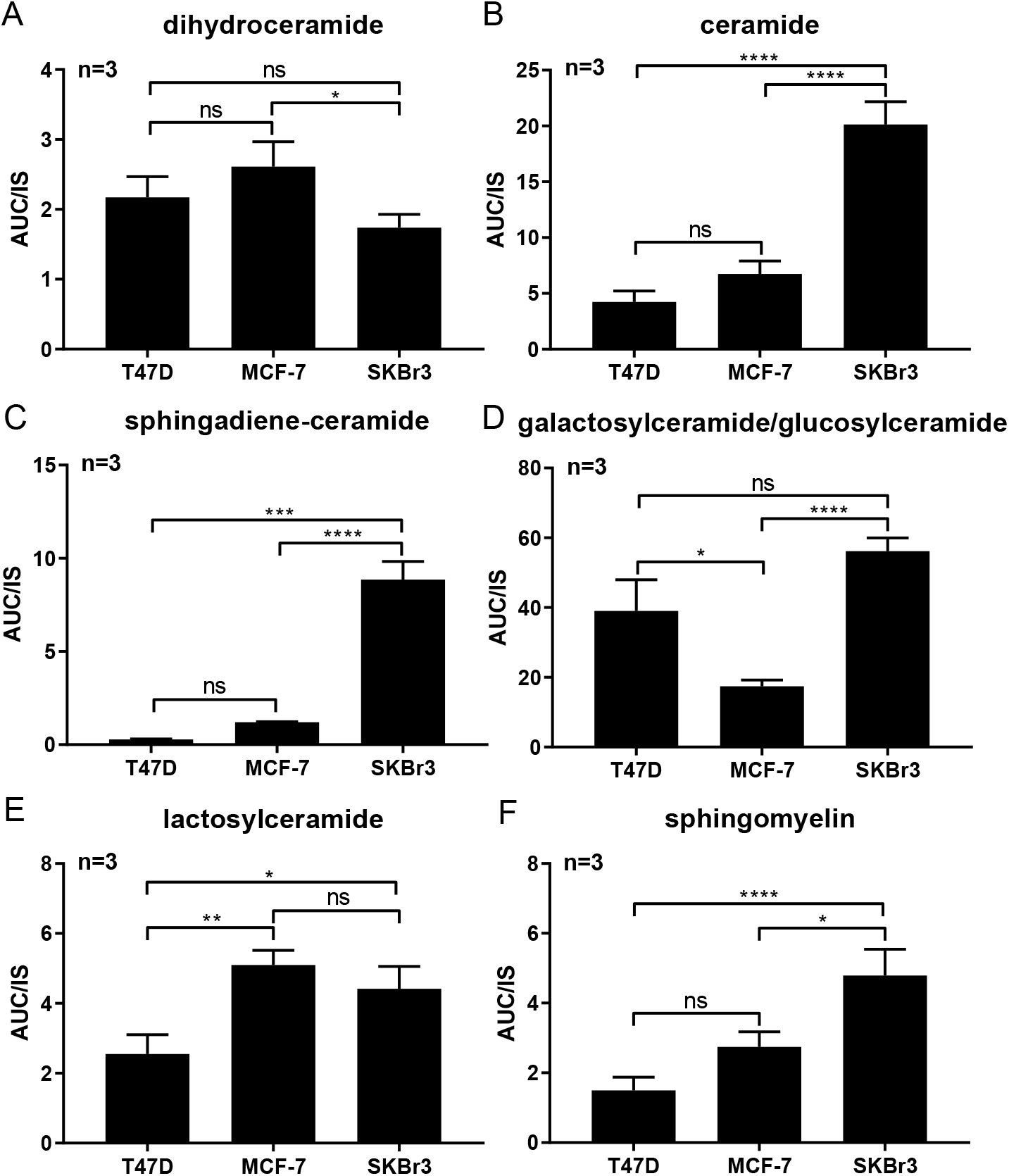
Sphingolipid species in breast cancer cells identified by LC-HRMS analysis. **A** Total *dihydroceramide* (dhCer) levels. Total of the following analytes: Cer dl8:0/22:0, Cer d18:0/24:1, Cer d18:0/24:1. B Total ceramide levels. Total of the following analytes: Cer d18:1/16:0, Cer d18:1/18:0, Cer d18:1/22:0, Cer d18:1/22:1, Cer d18:1/23:0, Cer d18:1/24:O, Cer d18:1/24:1. **C** Total sphingadiene-ceramide levels. Total of the following analytes: Cer d18:2/22:0, Cer d18:2/24:0. **D** Total *galactosylceramide* (GalCer)/*glucosylceramide* (GlcCer) levels. Total of the following analytes: HexCer d18:1/16:0, 24:0, 24:1. E Total *lactosylceramide* (LacCer) levels. Total of the following analytes: Hex2Cer d18:1/16:0, 24:1. **F** Total *sphingomyelin* (SM) levels. Total of the following analytes: SM 30:1, 32:1, 32:2, 33:1, 34:0, 34:1, 34:2, 36:1, 36:2, 36:3, 37:1, 38:1, 38:2, 40:1, 40:2, 40:3, 41:1, 41:2, 42:1, 42:2, 42:3, 43:1, 43:2. The identified single analytes are displayed in a heatmap in **supplemental data 1.** Data are presented as a mean of n=3 ± SEM. Tukey’s multiple comparison test. *\*p≤0.05, **p≤0.01, ***p≤0.001, ****p≤0.0001. Area under curve* (AUC), *internal standard* (IS), *not significant* (ns).

### 2.4. ER-and GPER1-dependent non-sphingolipid levels

LC-HRMS analysis revealed GPER1- and ER-dependent alterations of non-sphingolipid levels. Cholesterol levels are the lowest in T47D and the highest in SKBr3 cells **(Figure 3A).** Total sterol ester concentration is the lowest in MCF-7 cells and the highest in SKBr3 cells **(Figure 3B** and **supplemental data 2A).** By far the most altered lipid concentration between the three cell lines is the ether lipid concentration. SKBr3 cells exhibit a 140-fold increased ether lipid level compared to ER + cell lines **(Figure 3C** and **supplemental data 2B).** *Diglycerides* (DG) follow the same trend as cholesterol with T47D cells having the lowest and SKBr3 cells the highest concentration **(Figure 3D** and **supplemental data 2C).** In contrast, *triglyceride* (TG) content is low in MCF-7 cells as compared to T47D and SKBr3 cells **(Figure 3E** and **supplemental data 2D).** Glycerophospholipids appear most abundant in MCF-7 cells **(Figure 3F** and **supplemental data 2E),** whereas lyso-glycerophospholipids are the lowest in MCF-7 cells **(Figure 3G** and **supplemental data 2F).** Lyso-glycerophospholipids are statistically increased in SKBr3 cells as compared to T47D cells **(Figure 3G** and **supplemental data 2F).** Acylcarnitine levels are significantly decreased in SKBr3 cells compared to MCF-7 cells, whereas no difference compared to T47D cells could be detected **(Figure 3H** and **supplemental data 2G).** Treatment with the GPER1 antagonist G15 leads to significantly reduced levels of sterol ester and ether lipids in SKBr3 cells **(supplemental data 2H).** In contrast, G15 stimulation leads to significantly increased levels of cholesterol, diglyceride, glycerophospholipids and acylcarnitines **(supplemental data 2H).** SKBr3 cells exhibit increased sphingolipid levels as compared to the other cell lines, as well as increased levels of several non-sphingolipids such as cholesterol, DGs and ether lipids. This indicates that sphingolipid and non-sphingolipid pathways might be co-regulated in SKBr3 cells. In addition, treatment with the GPER1 antagonist G15 indicate GPER1-dependent co-regulation of sphingolipids and non-sphingolipids in GPER1 + breast cancer cells.

**Figure 3:**
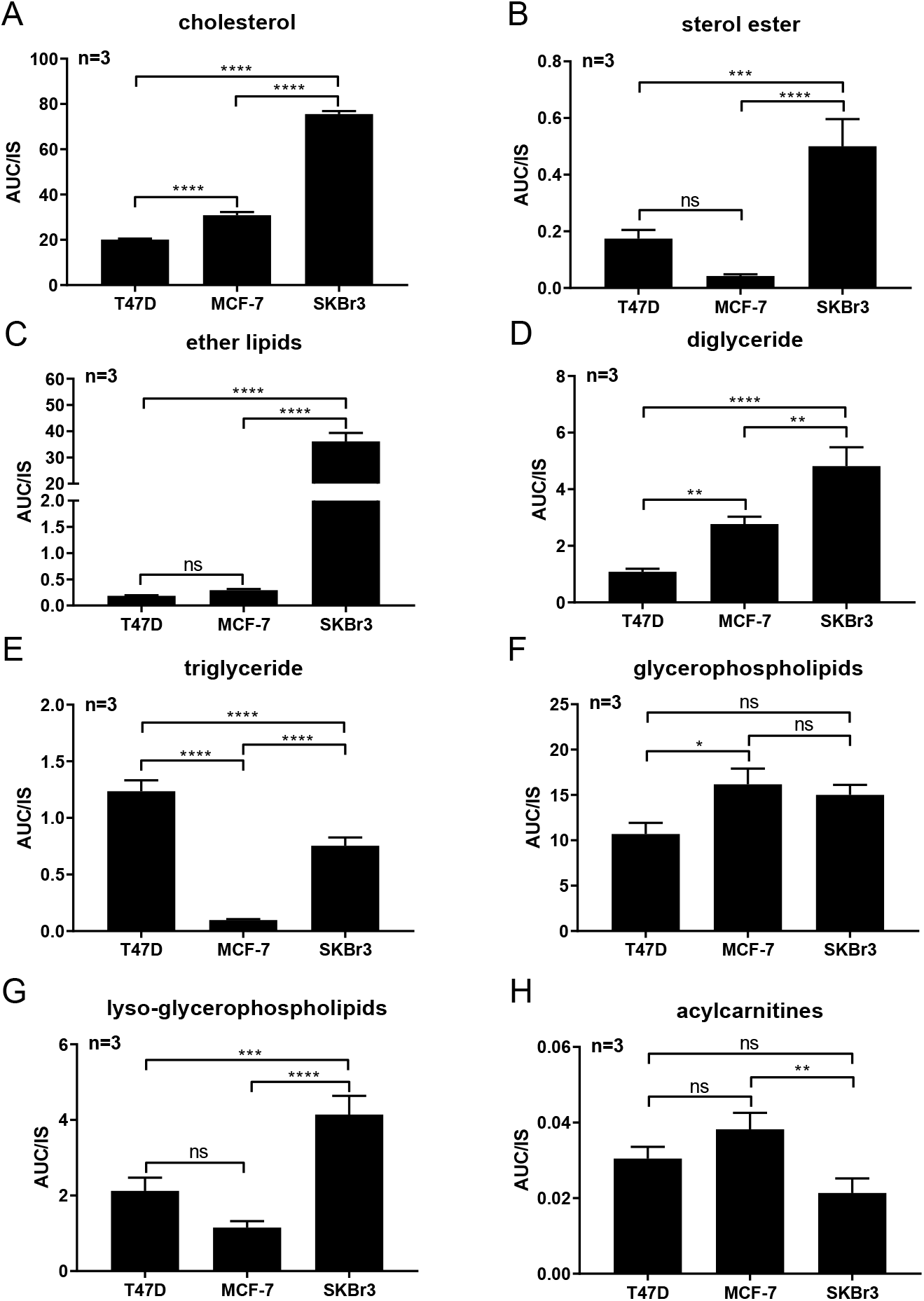
Non-sphingolipid species in breast cancer cells identified by LC-HRMS analysis. **A** Cholesterol (ST 27:1_OH). B Total sterol ester levels. Total of the following analytes: SE 27:1/14:1, 16:1, 17:1, 18:1, 18:2, 20:3, 20:4, 22:6, 24:1. **C** Total ether lipid levels. Total of the following analytes: LPC 0-16:0, 16:1, 18:0, 18:1, PC 0-16:0_16:0, 18:2, 20:4, PC O-16:1_16:0, 18:1, 18:2, 20:4, PC 0-l8:1_20:4, PC 0-34:1, PE O-16:1_18:2, 20:4, PE 0-18:1_18:1, 18:2, 20:4, LPE 0-16:1, 18:1. **D** Total *diglyceride* (DG) levels. Total of the following analytes: DG 32:1, 32:2, 34:1, 34:2, 34:3, 36:1, 36:2, 36:3, 38:2, 38:5. E Total *triglyceride* (TG) levels. Total of the following analytes: TG 42:1, 42:2, 43:0, 44:0, 44:1, 44:2, 46:0, 46:1, 46:2, 46:3, 48:0, 48:1, 48:2, 48:3, 48:4, 50:1, 50:2, 50:3, 50:4, 52:1, 52:2, 52:3, 52:4, 52:5, 54:1, 54:2, 54:3, 54:4, 54:5, 54:6, 56:1, 56:2, 56:3, 56:4, 56:6, 58:1, 58:2, 58:3, 58:4, 58:6. F Total glycerophospholipids levels. Total of the following analytes: PE 32:1, 34:1, 34:2, 34:3, 36:1, 36:2, 36:3, 36:4, 36:5, 38:4, 38:5, 38:6, 40:6, 40:7, PG 34:1, 36:2, PI 32:1, 34:1, 34:2, 36:1, 36:2, 36:4, 38:4, 38:6, 40:5, 40:6, PS 32:1, 34:1, 34:2, 36:1, 36:2, 38:2, 38:3, PC 30:0, 30:1, 30:2, 32:0, 32:1, 32:2, 33:2, 34:0, 34:1, 34:2, 34:3, 34:4, 34:5, 36:1, 36:2, 36:3, 36:4, 36:5, 38:2, 38:3, 38:4, 38:6, 38:7, 40:6, 40:7. G Total lyso-glycerophospholipid levels. Total of the following analytes: LPE 16:0, 18:0, LPG 16:0, 18:1, 18:2, LPI 16:0, 18:0, 18:2, 20:3, 20:4, LPS 18:0, 18:1, LPC 14:0, 15:0, 16:0, 17:0, 18:0, 18:3, 20:0, 20:1, 20:3, 22:0, 24:0. H Total acylcarnitine levels. Total of the following analytes: acylcarnitine 14:1, 16:0, 18:0, 18:1. The identified single analytes are displayed in heatmaps in **supplemental data 2.** Data are presented as a mean of *n=3* ± SEM. Tukey’s multiple comparison test. *\*p≤0.05, **p≤0.01, ***p≤0.001, ****p≤0.0001. Area under curve* (AUC), *internal standard* (IS), *not significant* (ns).

### 2.5. Effect of ER and GPER1 on cell proliferation

There are differences in morphology between the three cell lines such that T47D cells are the smallest in size, MCF-7 cells are larger and SKBr3 cells are the largest **(Figure 4A).** The results of the proliferation assay show that MCF-7 cells proliferate faster than SKBr3 cells, which in turn proliferate faster than T47D cells **(Figure 4B).** Since the three breast cancer cell lines differ in their ER and GPER1 status and in their sphingolipid and non-sphingolipid expression pattern, it is possible that cell size and proliferation is affected by this.

**Figure 4:**
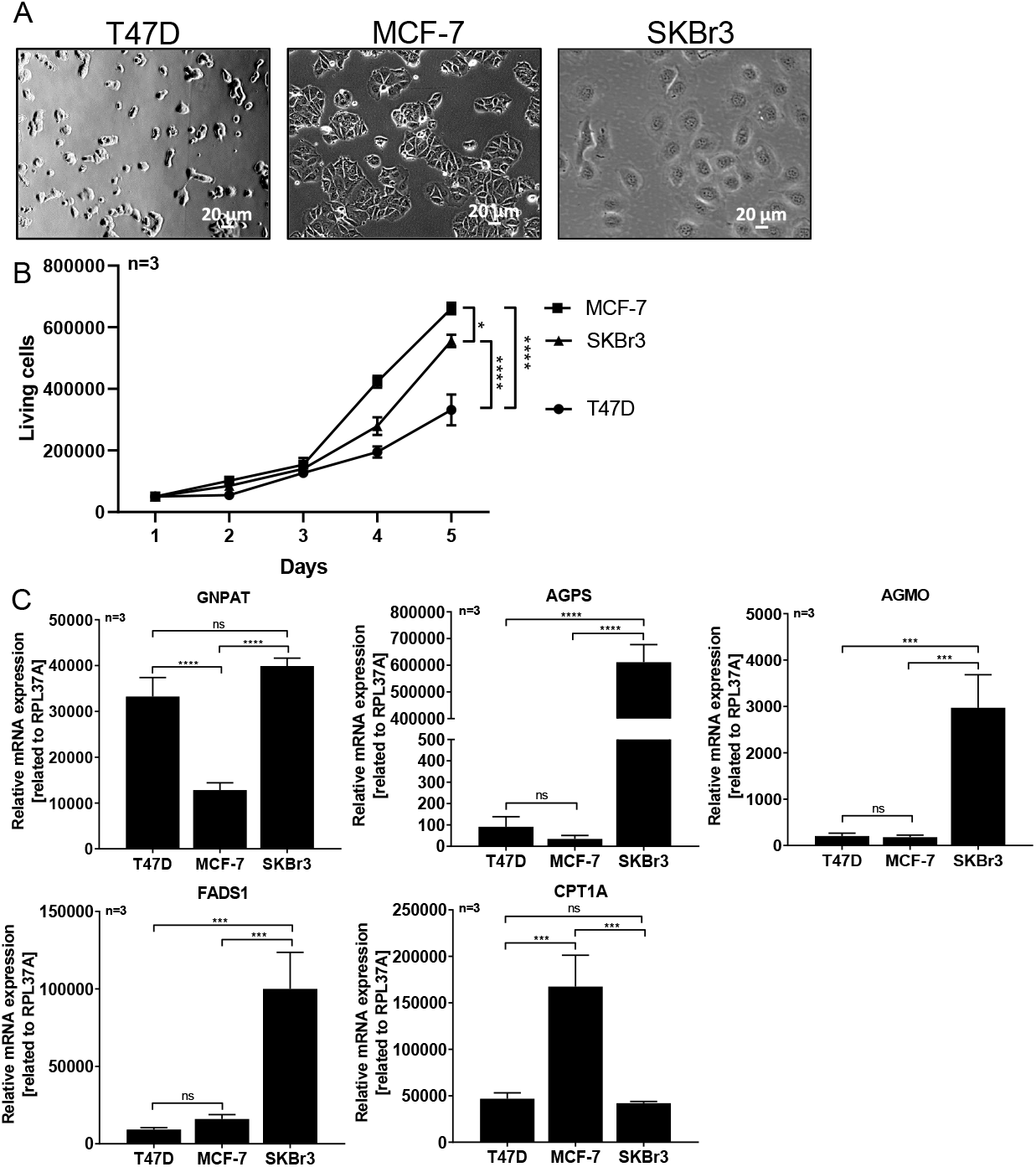
Morphology, proliferation and ether lipid metabolizing enzyme mRNA expression analysis by qRT-PCR of MCF-7, T47D and SKBr3 cells. **A** Light microscopy images of breast cancer cells with differing ER and GPER1 status. A representative image of each breast cancer cell line is displayed. B Living cell number on day 1, 2, 3, 4 and 5. Data are presented as a mean of *n=3 ±* SEM. Tukey’s multiple comparison test. **C** *Glyceronephosphate O-acyltransferase* (GNPAT), *alkylglycerone phosphate synthase* (AGPS), *alkylglycerol monooxygenase* (AGMO), *fatty acid desaturase 1* (FADS1) and *carnitine palmitoyltransferase 1A* (CPT1A) mRNA expression related to the housekeeping gene RPL37A. Data are presented as a mean of n=3 ± SEM. Tukey’s multiple comparison test. *\*p≤0.05, **p≤0.01, ***p≤0.001, ****p≤0.0001*.

### 2.6. GPER1-dependent ether lipid metabolizing enzymes mRNA expression

Since a strong GPER1-dependent increase of ether lipid level could be detected, we analyzed several enzymes involved in the ether lipid metabolism on mRNA level by qRT-PCR. *Glyceronephosphate O-acyltransferase* (GNPAT) is a crucial enzyme in ether lipid synthesis and is significantly lowered in MCF-7 cells compared to the other two cell lines **(Figure 4C).** Another key enzyme in ether lipid synthesis is *alkylglycerone phosphate synthase* (AGPS). AGPS is highly increased in SKBr3 cells, which fits the finding of strongly increased ether lipid concentrations in SKBr3 cells **(Figure 4C).** The *alkylglycerol monooxygenase* (AGMO) cleaves the O-alkyl bond of ether lipids leading to ether lipid degradation. Surprisingly, SKBr3 cells, which exhibit a high amount of ether lipids, also show strongly increased AGMO mRNA expression **(Figure 4C).** This might indicate a GPER1-dependent mRNA expression regulation of AGMO. In addition, *fatty acid desaturase 1* (FADS1) mRNA expression is strongly increased in SKBr3 cells **(Figure 4C).** *Carnitine palmitoyltransferase 1A* (CPT1A) mRNA expression in SKBr3 cells is significantly reduced compared to MCF-7 cells, whereas no difference could be detected as compared to T47D, which verifies the LC-HRMS analysis results for acylcarnitine levels **(Figure 4C).** When normalizing LC-HRMS analysis results using *median peak ratio* (MPR) as compared to one *internal standard* (IS) per lipid class, the distribution of the reading points does not change **(Figure 5** and **supplemental data 3),** whereby effects ascribable to cell size are excluded. In summary, our data indicate high ether lipid turnover in GPER1 +, but ER - cells (SKBr3) and a putative GPER1-dependent regulation of the metabolizing enzymes.

**Figure 5:**
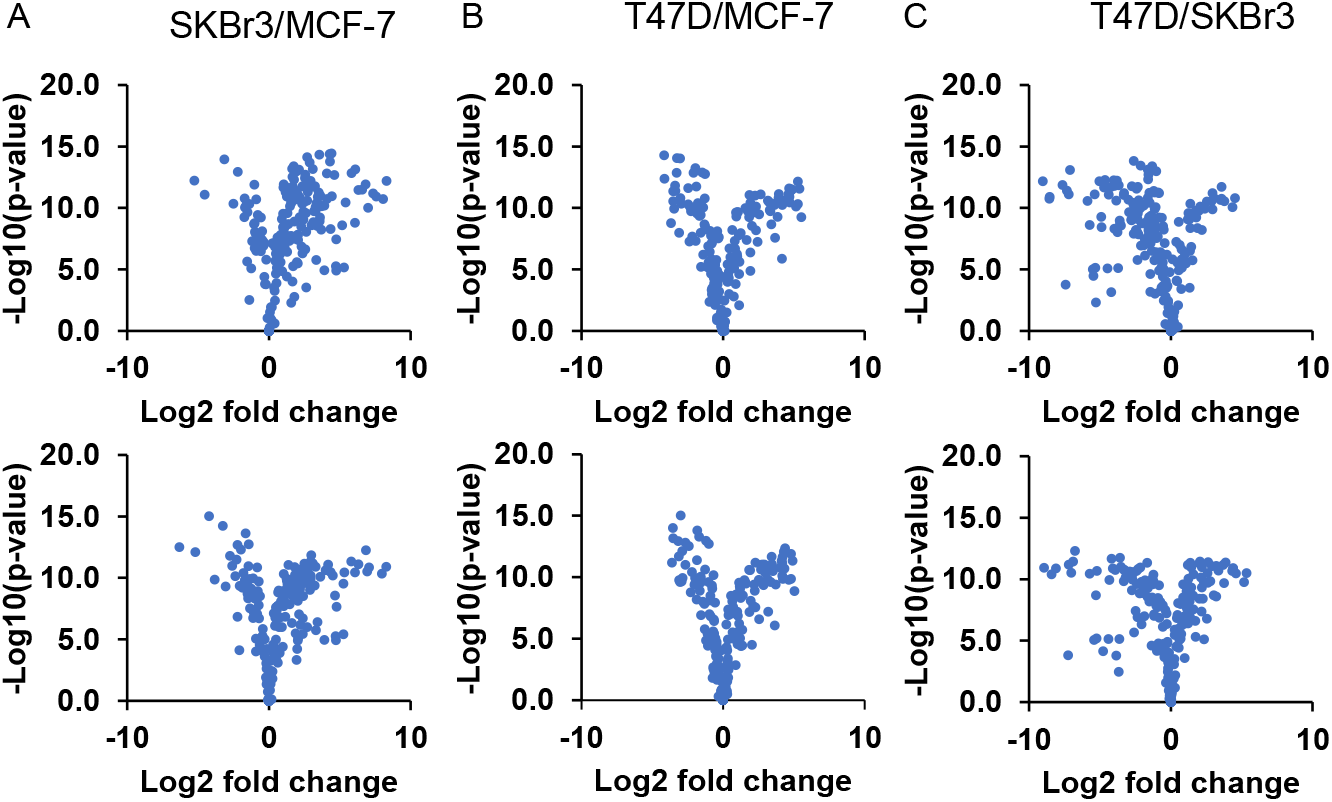
*Internal Standard* (IS) normalized LC-HRMS data (upper row) and *median peak ration* (MPR) normalized LC-HRMS data (lower row). A SKBr3/MCF-7. B T47D/MCF-7. C SKBr3/T47D.

## 3. Materials and Methods

### 3.1. Cell lines

MCF-7, T47D and SKBr3 cells were purchased from the Health Protection Agency (European Collection of Cell Cultures, ECACC, Salisbury, UK) and were cultured in phenol-red free *Dulbecco’s Modified Eagle’s Medium* (DMEM) supplemented with 5 % charcoaled *fetal bovine serum* (FBS), 1 % sodium pyruvate and 1 % GlutaMAX. They were maintained in a humidified, 5 % CO_2_ supplied atmosphere incubator at 37 °C.

### 3.2. Quantitative real-time-PCR (qRT-PCR)

Total RNA was isolated using RNeasy Mini Kit (QIAGEN, Hilden, Germany). cDNA was synthesized from 300 ng total RNA using VERSO™ cDNA Synthesis Kit (Thermo Fisher Scientific, ABgene, Epsom, UK). Gene specific PCR products were assayed using Maxima EvaGreen qPCR Master Mix on a 7500fast quantitative PCR system (TaqMan®, Life Technologies, Darmstadt, Germany). Relative gene expression was determined using the ΔCT method, normalizing relative values to the expression level of *60S ribosomal protein L37a* (RPL37A) as a housekeeping gene. It is shown that RPL37A is the optimal single reference gene when normalizing gene expression in meningiomas and control tissue (Pfister et al., 2011). Furthermore, Maltseva et al. showed that RPL37A has similar high expression stability values compared to the other genes such as ACTB or RPS23 in breast cancer cells (Maltseva et al., 2013). Accordingly, RPL37A is a suitable housekeeper gene for our study. The primer mix for *sphingomyelin synthase 1* und 2 (SMS1 and SMS2) and *galactosylceramide synthase* (GalCerS) detection were purchased from GeneCopeia (Rockville, USA) and the GPER1-primermix from Realtimeprimers (Pennsylvania, USA).

**Table 1.**
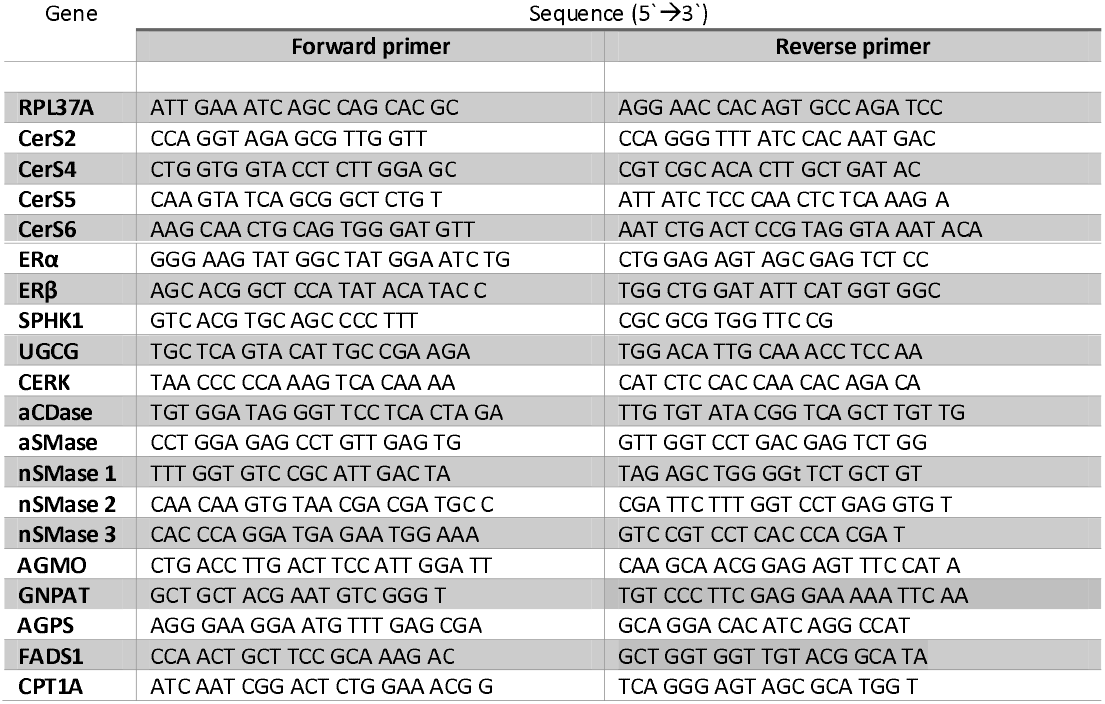
qRT-PCR primer sequences.

### 3.3. Lipidomics analysis

#### 3.3.1. Materials

Water, isopropanol, methanol (LC-MS grade) and *methyl-tert-butyl-ether* (MTBE, HPLC-grade) were purchased from Carl Roth (Karlsruhe, Germany). Ammonium formate (for mass spectrometry, ≥ 99.0 %) and Sulfinpyrazon (100 *%) were* purchased from Sigma-Aldrich (Munich, Germany) and acetonitrile (ULC-MS grade) from Biosolve B. V. (Valkenswaard, Netherlands). APCI positive calibration solutions was obtained from Sciex (Darmstadt, Germany) and formic acid (98-100 %) from AppliChem (Darmstadt, Germany). All internal standards were purchased from Avanti Polar Lipids (Alabaster, AL, USA).

#### 3.3.2. Liquid chromatography time-of-flight mass spectrometry (LC-HRMS)

Three experiments with three replicates each were performed. The sample processing and LC-HRMS measurement was carried out as described previously (Hahnefeld et al., 2020). Approximately 5 x 10^5^ cells in 150 μL PBS were thawed in the fridge for 30 min before extraction with 150 μL internal standards in methanol and 500 μL MTBE. After centrifugation at 20,000 g for 5 min, the upper organic phase was transferred and the aqueous phase was reextracted using 200 μL of MTBE: methanol: water (10:3:2.5, v/v/v). The combined organic phases were split for measurement in positive and negative ionization mode, dried at 45 °C under a nitrogen stream and stored at −20 °C pending analysis. The samples were dissolved in 120 μL methanol before analysis. For quality control, the different cell lines were pooled with two injections at the beginning and the end of each run and one after every 10th sample. The LC-MS measurement was carried out on a Shimadzu Nexera-X2 (Shimadzu Corporation, Kyoto, Japan) with a Zorbax RRHD Eclipse Plus C8 1.8 μm 50×2.1 mm ID column (Agilent, Waldbronn, Germany) coupled to TripleTOF 6600 (Sciex, Darmstadt, Germany) with electrospray ionization operating in positive and negative ionization mode. A mass range from 100 to 1000 m/z was scanned together with data-dependent acquisition for improved identification with a mass error of ± 5 ppm. The data acquisition was performed using Analyst TF v1.71. Compound identification and semi-targeted analysis was achieved with MasterView v1.1 and MultiQuant v3.02 software as described previously (Hahnefeld et al., 2020). The software was obtained from Sciex (Darmstadt, Germany).

#### 3.3.3. Sample Normalization

LC-HRMS analysis results were compared after normalization once with one *internal standard* (IS) per lipid class and once with *median peak ratio* (MPR) as calculated by MarkerView v1.1 software. For MPR calculation a selected reference sample, usually the first QC sample, is used to generate a list of all peaks with a minimum peak area of 1 % of the largest signal. For each peak in the list the peak area is divided by the peak area of the reference sample and subsequently the median of the area ratios is calculated. If no peak appears in the analyzed sample, the area ratio is set to the value 1. Accordingly, a normalization factor for each sample was generated and the values of all analytes were multiplied by this normalization factor. The results normalized with one internal standard per lipid class were used for further analysis.

### 3.4. Proliferation assay

For quantitative proliferation assays, cells were seeded at a density of 5 x 10^4^ cells/well of a 6-well plate (cell culture multiwell plate, 6 well, clear, sterile, (Greiner AG, Kremsmünster, Austria)). Since the cells were not 100 % confluent following 5 days of culture, the media was not replenished during the 5 days of culture. Cells were harvested at day 1, 2, 3, 4, 5 and living cell number was counted using a Neubauer counting chamber and trypan blue (labels dead cells exclusively).

### 3.5. Statistical analysis

Data are presented as mean ± *standard error of the mean* (SEM). Statistical analyses were performed with GraphPad Prism 7 software. Significant differences (p < 0.05) between groups were assessed by Tukey’s multiple comparison test as indicated in the figure descriptions.

## 4. Discussion

With our LC-HRMS analysis approach we were able to show co-expression of sphingolipids and non-sphingolipids in breast cancer cells. Importantly, GPER1 might be involved in regulating this co-expression of different lipid species.

Our analysis identified a distinct mRNA expression pattern in both anabolic and catabolic sphingolipid metabolizing enzymes among the three breast cancer cell lines tested. Low CerS5 and CerS6 mRNA expression (responsible for the production of C16 to C20 ceramides) in SKBr3 cells (GPER1 +, ER -) is in line with results we published previously showing decreased CerS5 and CerS6 mRNA (production of C16 ceramides) level in MCF-7 cells following stable GPER1 plasmid transfection (MCF-7/GPER1) (Wegner et al., 2019). However, following GPER1 overexpression in MCF-7 cells, CerS2 mRNA (production of C20-26 ceramides) level is increased and CerS4 mRNA level unchanged (Wegner et al., 2019), whereas SKBr3 cells exhibit lower CerS2 and higher CerS4 mRNA levels compared to MCF-7 wild type cells. This indicates that regulation of CerS2 and CerS4 is regulated by both GPER1 and ER and that the lack of ERα and β in SKBr3 cells contributes to the differences in SKBr3 and MCF-7/GPER1 cells. GPER1-dependent SPHK1 activity is shown (reviewed in (Sukocheva and Wadham, 2014)) and is in line with our data, which reveal high SPHK1 mRNA levels in GPER1 + cells (SKBr3). Sukocheva et al. have shown estrogen-dependent SPHK1 activation that leads to *sphingosine 1-phosphate* (S1P) release, accordingly to Edg-3 activation and results in *enhanced growth factor receptor* (EGFR) transactivation (Sukocheva et al., 2006). Our previous data show that GPER1 regulates CerS transcription ligand-independently (Wegner et al., 2014). Also, since phenol-red is known to mediate estrogen-like mechanisms (Wesierska-Gadek et al., 2007), we used phenol-red free media and 5 % charcoaled FBS in our current study. This could indicate that in addition to ligand-dependent mechanisms, ligand-independent mechanism mediated by ERs could also contribute to SPHK1 regulation. Since SPHK1 is a marker for poor prognosis in breast cancer patients (reviewed in (Sukocheva and Wadham, 2014)) it would be intriguing to investigate GPER1-dependent mechanisms of SPHK1 regulation. No distinct differences on SMS1 and SMS2 mRNA expression levels between the three cell lines were observed indicating that SMS are constitutively expressed. However, we detected SMS1 and SMS2 mRNA expression increase following stable GPER1 overexpression in MCF-7 cells (Wegner et al., 2019). This indicates a putative interaction of GPER1 and ERα and or β. Moro et al. showed suppression of CERK expression in cancer cells (Moro et al., 2018). The breast cancer cell line SKBr3 exhibits strongly increased CERK mRNA expression. This indicates that high CERK expression might not be a prognostic marker for cancer in general, but rather identifies a less aggressive type of cancer. Interestingly, the product of CERK activity, *ceramide 1-phosphate* (C1P) is a known inducer of inflammatory processes in cancer cells, which could contribute to the activation of pro-cancerous signaling pathways (reviewed in (Dei Cas and Ghidoni, 2018, Arana et al., 2010)). Indeed, we measured increased FADS1 mRNA expression in SKBr3 cells. FADS1 catalyzes the final step in arachidonic acid synthesis. However, Owczarek et al. showed that GalCerS, which is also highly expressed in SKBr3 cells, functions as a pro-tumorigenic protein by inhibiting apoptosis (Owczarek et al., 2013). This is contradictory to the finding that SKBr3 cells are less aggressive, however, we have already shown that UGCG overexpression in MCF-7 cells leads to increased glutamine uptake (Schomel et al., 2019). This results in reinforced oxidative stress response and fueled *tricarboxylic acid* (TCA) cycle, which is accompanied by increased cell proliferation (Schomel et al., 2019). Since MCF-7 cells exhibit a p53 wildtype while T47D (Lim et al., 2009) and SKBr3 cells (Garufi et al., 2016) display a p53 mutant status and a high UGCG mRNA expression, it is likely that UGCG overexpression is connected to p53 signaling pathway inhibition. In line with this, Liu et al. showed that UGCG suppression restores apoptosis mediated by p53 in mutant p53 cancer cells (Liu et al., 2011). However, MCF-7 cells exhibit the most increased aCDase mRNA expression as well as increased proliferation. This fits the finding of Lucki et al. showing aCDase driven increased proliferation of MCF-7 cells (Lucki and Sewer, 2011). Usually, aCDase overexpression in breast cancer patients correlates with a better prognosis (Ruckhaberle et al., 2009, Sanger et al., 2015).

Overall, levels of anabolic enzyme mRNA expression are more distinct than the mRNA levels of catabolic enzymes in the analyzed cell lines. Increased anabolic sphingolipid enzyme mRNA expression levels are indicated by an increase in total ceramide, sphingadiene-ceramide, GalCer/GlcCer and SM levels in SKBr3 cells. Each sample contained the same cell number, but SKBr3 cells are larger in size than T47D and MCF-7 cells. We assumed that the total metabolite concentration might correlate with cell size and therefore differences in cell size are compensated by normalization with MPR (Muschet et al., 2016). Normalization with one IS per lipid class cannot compensate for unequal cell size, but often improves measurement deviations. The volcano plots show that the differences between lipid levels are not ascribable to cell size, whereas the proliferation differs between the cell lines. However, since dhCer levels are decreased and ceramide levels strongly increased in SKBr3 cells, it is likely that the *dihydroceramide desaturase 1* (DES1) activity is elevated. Interestingly, SKBr3 cells exhibit strongly increased sphingadiene-ceramide concentration. Information about the biological function of sphingadiene-ceramides is limited. Sphingadiene-ceramides inhibit the *phosphoinositide 3-kinase* (PI3K)/Akt and Wnt signaling pathway leading to apoptosis (reviewed in (Hannun, 2015)).

Furthermore, we observed similar sphingolipid and non-sphingolipid expression patterns in SKBr3 cells indicating a co-regulation of these lipid species. Especially, ether lipids are strongly increased in SKBr3 cells, which is reversible by treatment with G15, a GPER1 antagonist. This is confirmed by strongly increased AGPS (generation of ether lipids) mRNA expression **(Summary Figure).** Interestingly, it is shown that aggressive cancer cells exhibit increased AGPS expression and ether lipid metabolism (Benjamin et al., 2013). Furthermore, Chen et al. postulated that plasma ether-linked *phosphocholine* (PC) species can be used as a biomarker for the diagnosis of breast cancer (Chen et al., 2016). One limitation of the study from Chen et al. is the lack of information about the ER and GPER1 status of the breast cancer. Our results do not confirm a general induction of ether lipid concentration in breast cancer cells, but indicate a GPER1-dependent regulation. Since SKBr3 cells are a non-aggressive cell line, GPER1-dependent increase of ether lipid synthesis does not state aggressiveness of breast cancer cells in general. However, AGPS is located in the peroxisome and is the rate-limiting enzyme in ether lipid synthesis (reviewed in (Dean and Lodhi, 2018)). Surprisingly, GNPAT, which is also located in the peroxisome and essential for ether lipid synthesis is only compared to MCF-7 cells significantly increased in SKBr3 cells. The reason for this could be the finding that GNPAT enzyme activity does not only require AGPS presence, but also depends on the integrity of channeling the substrate from GNPAT to AGPS, which is shown by Itzkovitz et al. (Itzkovitz et al., 2012). Therefore, GNPAT activity could be increased in SKBr3 cells leading to increased ether lipid synthesis without an increased GNPAT mRNA level. *Phosphocholine* (PC)-ether species could not be detected in T47D cells, but T47D cells exhibit a similar GNPAT mRNA expression as SKBr3 cells. GNPAT does not contribute to ether lipid production in T47D cells. Either GNPAT is less active in T47D cells or is expressed, because the enzyme also executes other tasks in T47D cells. AGMO, an ether lipid cleaving enzyme, also exhibits strongly increased mRNA levels in SKBr3 cells. This indicates that SKBr3 cells exhibit an accelerated ether lipid metabolism **(Summary Figure).** A crosstalk between ether lipids and sphingolipids has been shown in pathophysiological processes such as cancer and atopic dermatitis (reviewed in (Jimenez-Rojo and Riezman, 2019)). GlcCer levels in cancer have been shown to correlate to ether lipids through mechanisms that are linked to *mammalian target of rapamycin* (mTOR) signaling, which is activated in most tumors (Guri et al., 2017). Therefore, previous studies confirm our data showing that high ether lipid levels correlate with GSL levels in SKBr3 cells. Ether lipids can be covalently attached to proteins as components of *glycosylphosphatidylinositol* (GPI)-anchors. GPI-anchored proteins are linked to the membrane in the ER via the hydrophobic part of the glycolipid and are mostly delivered to the cell surface to execute diverse functions (reviewed in (Jimenez-Rojo and Riezman, 2019)). Typically GPI-anchored proteins are enriched in membrane microdomains (rafts) and these microdomains exhibit high sphingolipid and cholesterol levels (reviewed in (Kinoshita, 2016)). In addition to increased ether lipid concentration, SKBr3 cells exhibit elevated cholesterol levels. This indicates either increased raft formation or changed lipid composition of rafts as compared to the other cell lines. Ether lipids also affect membrane fluidity and cellular processes such as membrane fusion (reviewed in (Dean and Lodhi, 2018)). However, SKBr3 cells exhibit elevated DG levels. One possible mechanism leading to increased DG levels is GPER1-dependent phospholipase C activation, which results in DG production (reviewed in (Newton et al., 2016)). DGs function as second messengers in the cell by activating protein kinase C (PKC) and PKC function in the cell is manifold (reviewed in (Newton, 2018)). Interestingly, TG are the lowest in MCF-7 cells. Lofterød et al. showed that increased TG levels are connected to promoted tumor growth (Lofterod et al., 2018), which underlines the finding that SKBr3 cells have poor tumorigenic potential. Another interesting finding is the low CPT1A mRNA level in SKBr3 cells, which follows the same trend as total acylcarnitine level. Following production by CPT1A acylcarnitine is imported into mitochondria and used for β-oxidation in form of acyl-CoA (reviewed in (Schooneman et al., 2013)). The β-oxidation products *nicotinamide adenine dinucleotide* (NADH) and reduced *flavin adenine dinucleotide* (FADH_2_) are oxidized at the *oxidative phosphorylation* (OXPHOS) complexes I and II. This generates the mitochondrial membrane potential, which is essential for proper mitochondrial respiration function. The data indicate that SKBr3 cells execute less β-oxidation, which might lead to reduced mitochondrial respiration. We have shown that GPER1 overexpression in MCF-7 cells leads to decreased basal respiration and reduced glycolysis rate resulting in reduced cell proliferation (Wegner et al., 2019). However, acyl-CoA needed for ether lipid synthesis could be generated by increased de *novo* lipogenesis.

## 5. Conclusion

In conclusion, the results of our semi-targeted analysis show co-regulation of sphingolipids and non-sphingolipids in breast cancer cells with differing ER and GERP1 status. Especially, GPER1 seems to influence expression of sphingolipids and ether lipids. Importantly, this co-regulation might lead to a less tumorigenic potential. This finding might contribute to identification of novel potential therapeutic targets in breast cancer treatment.

## 6. Acknowledgements

This work was funded by the Deutsche Forschungsgemeinschaft (WE 5825/1-1 and WE 5825/2-1), the August Scheidel-Stiftung, the Heinrich und Fritz Riese-Stiftung, Johanna Quandt-Jubiläumsfond and the Paul und Ursula Klein-Stiftung. Support by the SFB 1039 is also gratefully acknowledged. The authors thank Ellen M. Olzomer for the linguistic revision of the manuscript.

## 8. Author approvals

All authors have seen and approved the manuscript. Furthermore, we ensure that the manuscript hasn’t been accepted or published elsewhere.

---

**Summary Figure:**
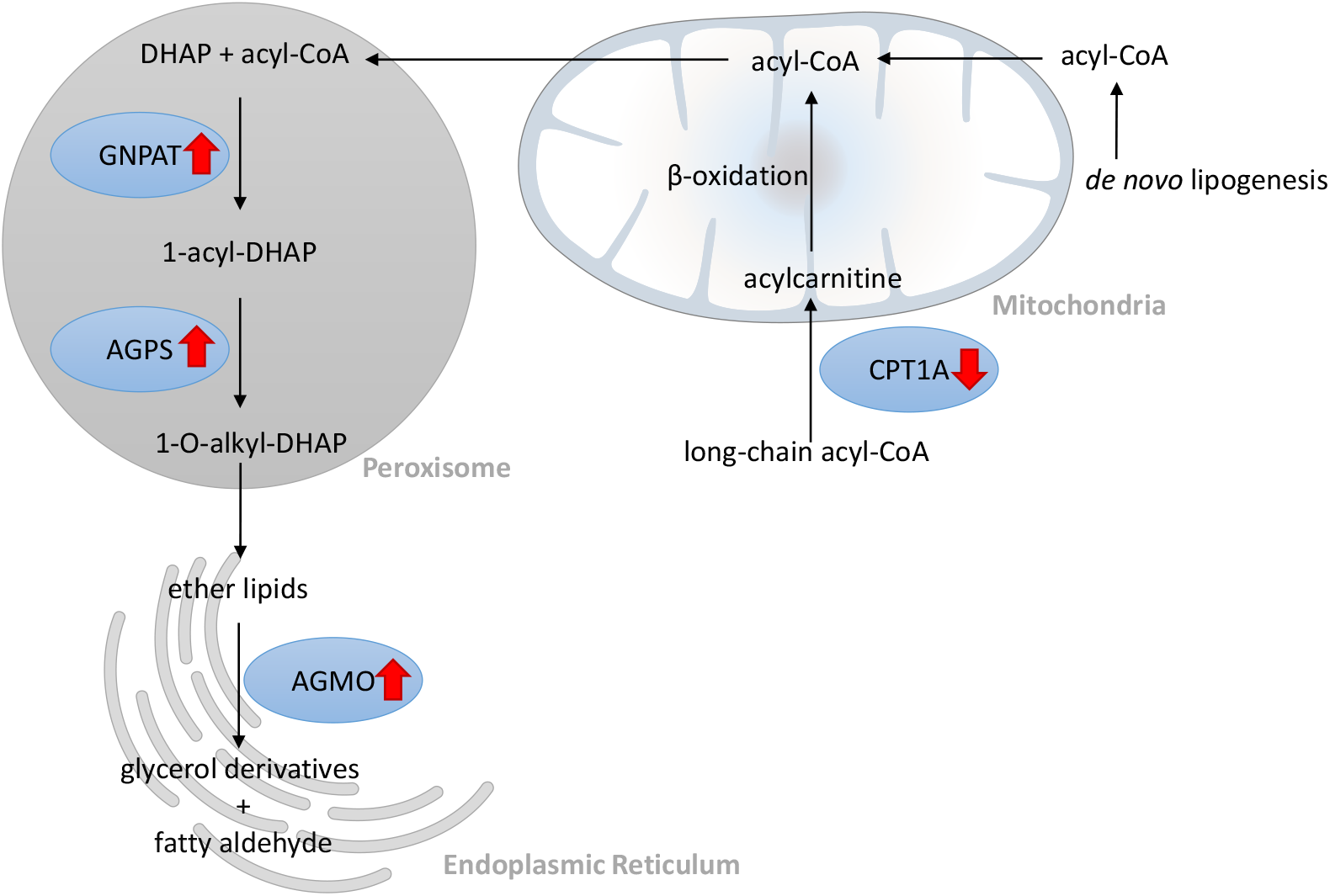
Summary of ether lipid metabolism alterations in GPER1 +, but ER - cells. The acyl CoA required for ether lipid synthesis in SKBr3 cells might be generated by *de novo* lipogenesis rather than β-oxidation. Both key enzymes of ether lipid synthesis (*glyceronephosphate O-acyltransferase* (GNPAT) and *alkylglycerone phosphate synthase* (AGPS)) are strongly increased in SKBr3 cells. Since ether lipid degrading *alkylglycerol monooxygenase* (AGMO) is also increased, accelerated ether lipid metabolism in GPER1 + (ER -) cells is assumed (red arrow = displays changes in mRNA expression level).

