## Supplemental for "Ether lipid and sphingolipid expression patterns are G-protein coupled estrogen receptor 1-dependently altered in breast cancer cells"

### Slide 1
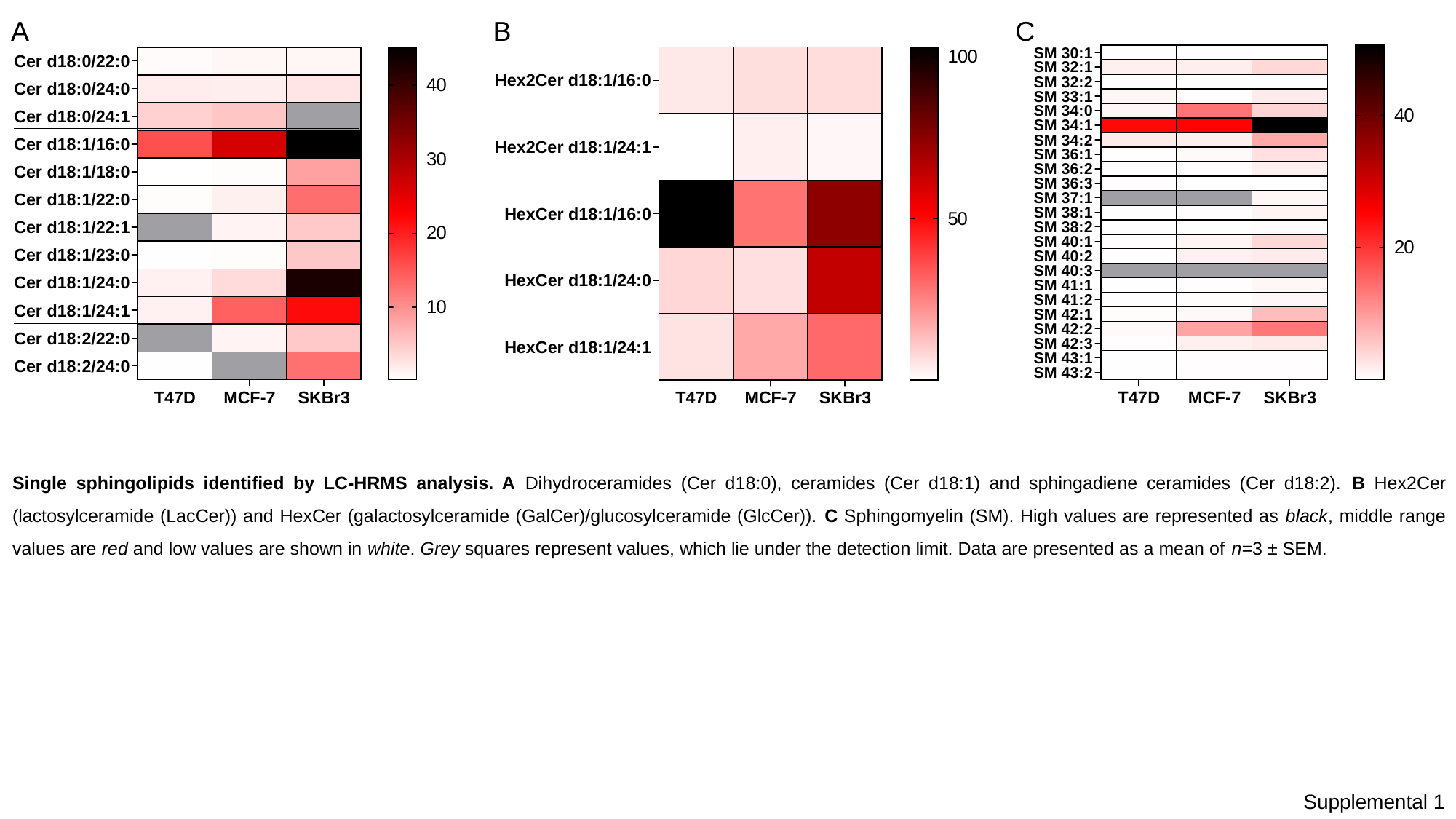

A
B
C
Single sphingolipids identified by LC-HRMS analysis. A Dihydroceramides (Cer d18:0), ceramides (Cer d18:1) and sphingadiene ceramides (Cer d18:2). B Hex2Cer (lactosylceramide (LacCer)) and HexCer (galactosylceramide (GalCer)/glucosylceramide (GlcCer)). C Sphingomyelin (SM). High values are represented as black, middle range values are red and low values are shown in white. Grey squares represent values, which lie under the detection limit. Data are presented as a mean of n=3 ± SEM.
Supplemental 1

### Slide 2
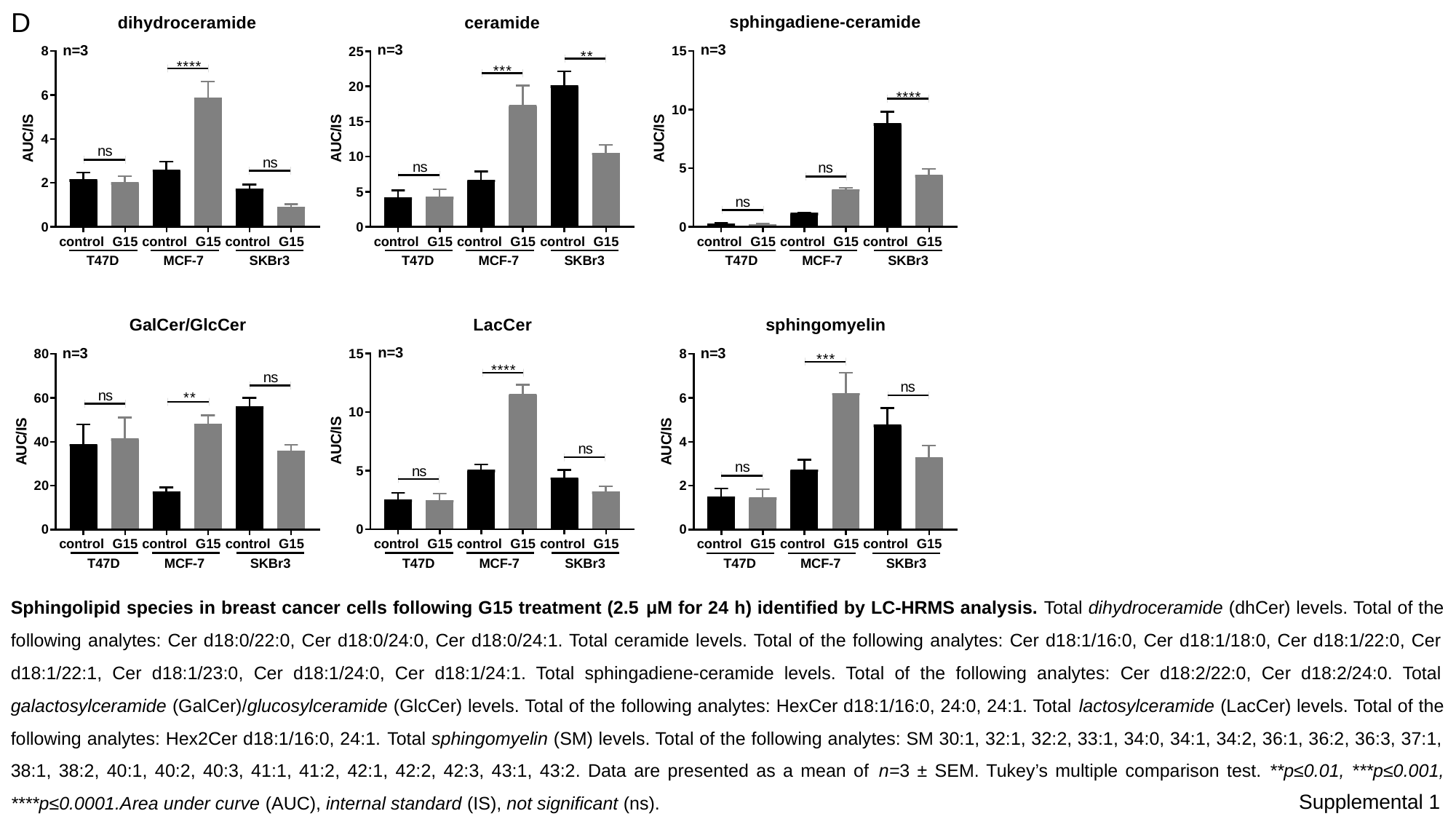

D
Sphingolipid species in breast cancer cells following G15 treatment (2.5 μM for 24 h) identified by LC-HRMS analysis. Total dihydroceramide (dhCer) levels. Total of the following analytes: Cer d18:0/22:0, Cer d18:0/24:0, Cer d18:0/24:1. Total ceramide levels. Total of the following analytes: Cer d18:1/16:0, Cer d18:1/18:0, Cer d18:1/22:0, Cer d18:1/22:1, Cer d18:1/23:0, Cer d18:1/24:0, Cer d18:1/24:1. Total sphingadiene-ceramide levels. Total of the following analytes: Cer d18:2/22:0, Cer d18:2/24:0. Total galactosylceramide (GalCer)/glucosylceramide (GlcCer) levels. Total of the following analytes: HexCer d18:1/16:0, 24:0, 24:1. Total lactosylceramide (LacCer) levels. Total of the following analytes: Hex2Cer d18:1/16:0, 24:1. Total sphingomyelin (SM) levels. Total of the following analytes: SM 30:1, 32:1, 32:2, 33:1, 34:0, 34:1, 34:2, 36:1, 36:2, 36:3, 37:1, 38:1, 38:2, 40:1, 40:2, 40:3, 41:1, 41:2, 42:1, 42:2, 42:3, 43:1, 43:2. Data are presented as a mean of n=3 ± SEM. Tukey’s multiple comparison test. **p≤0.01, ***p≤0.001, ****p≤0.0001.Area under curve (AUC), internal standard (IS), not significant (ns).
Supplemental 1

### Slide 3
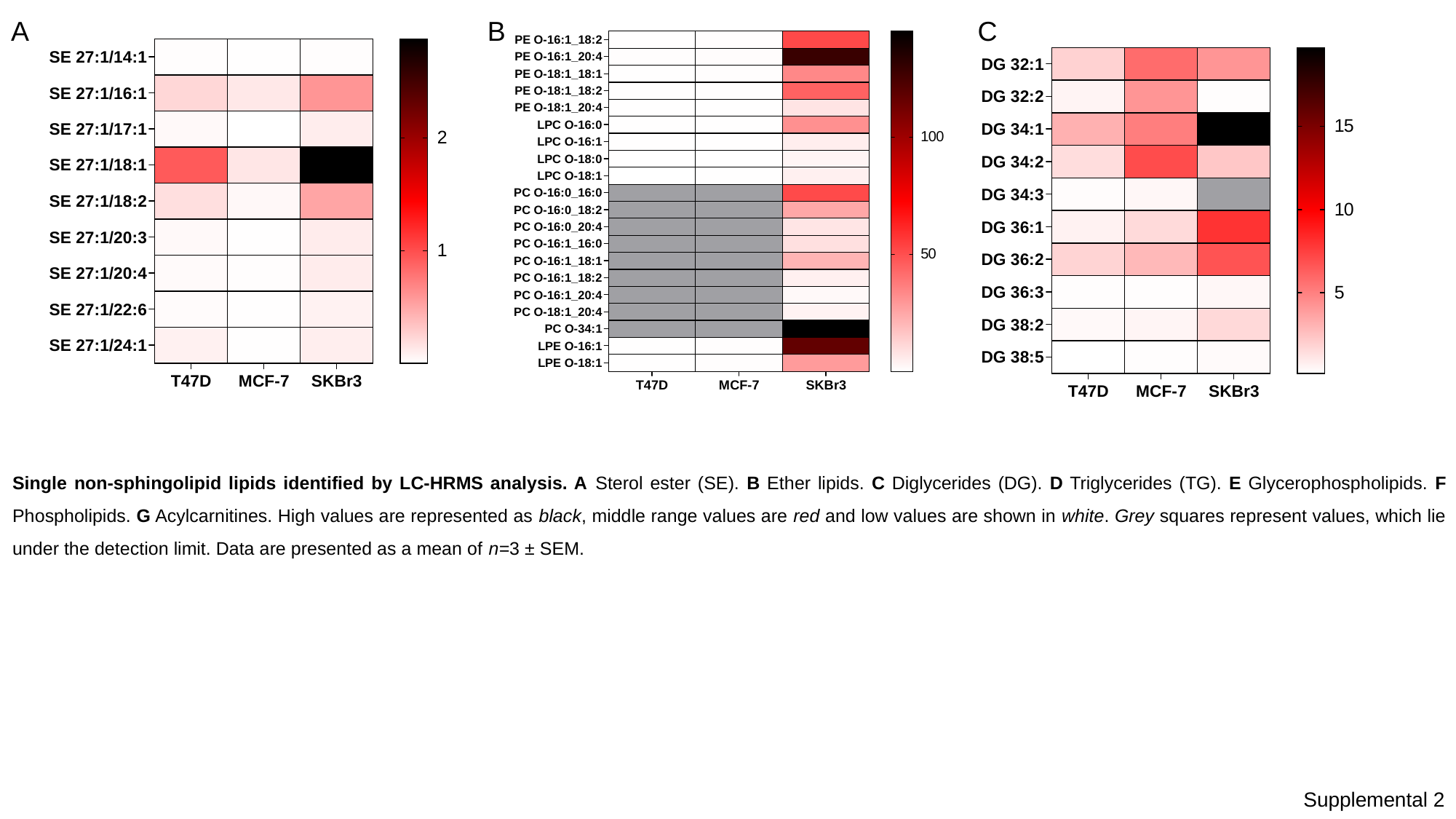

A
B
C
Single non-sphingolipid lipids identified by LC-HRMS analysis. A Sterol ester (SE). B Ether lipids. C Diglycerides (DG). D Triglycerides (TG). E Glycerophospholipids. F Phospholipids. G Acylcarnitines. High values are represented as black, middle range values are red and low values are shown in white. Grey squares represent values, which lie under the detection limit. Data are presented as a mean of n=3 ± SEM.
Supplemental 2

### Slide 4
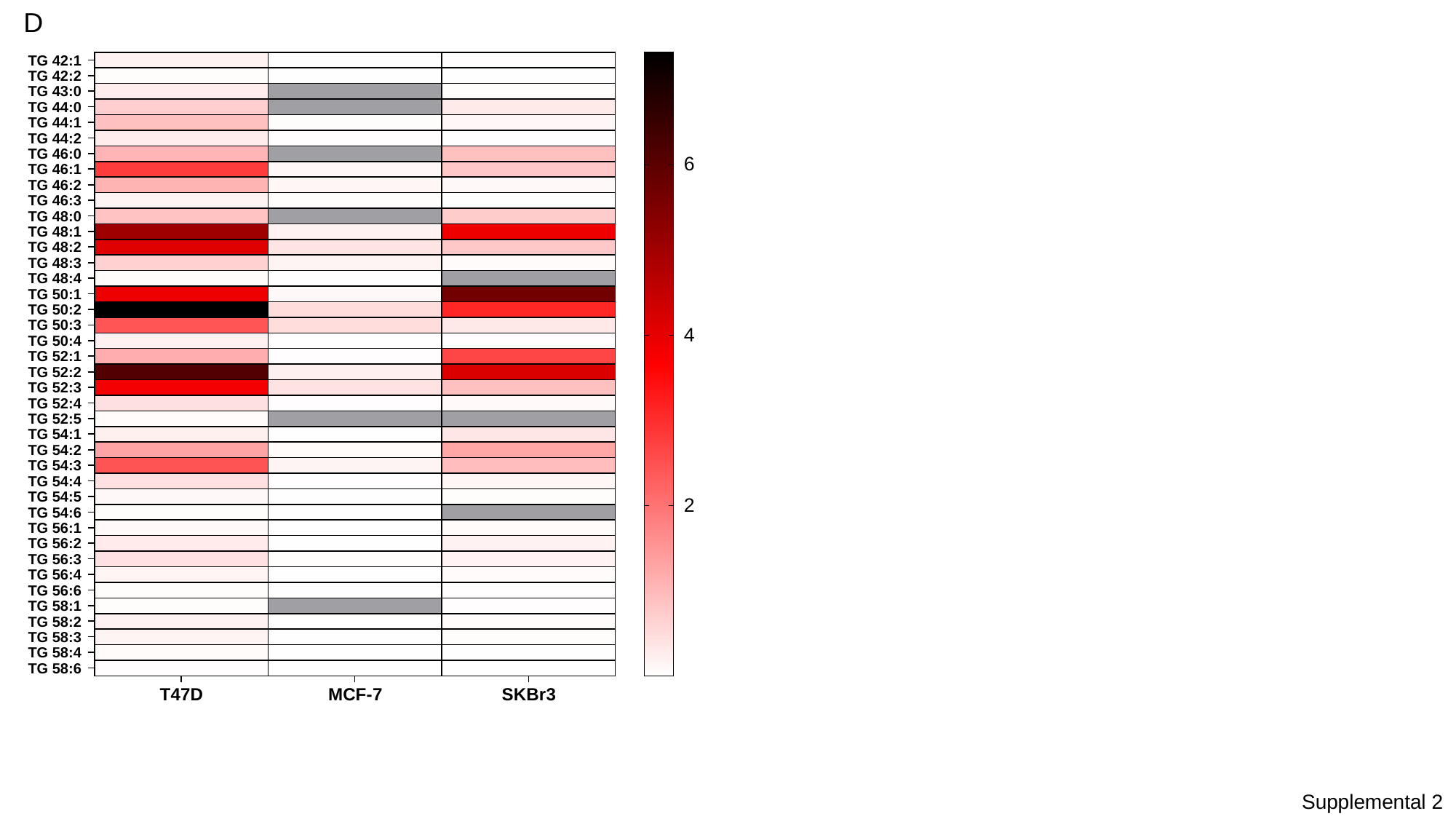

D
Supplemental 2

### Slide 5
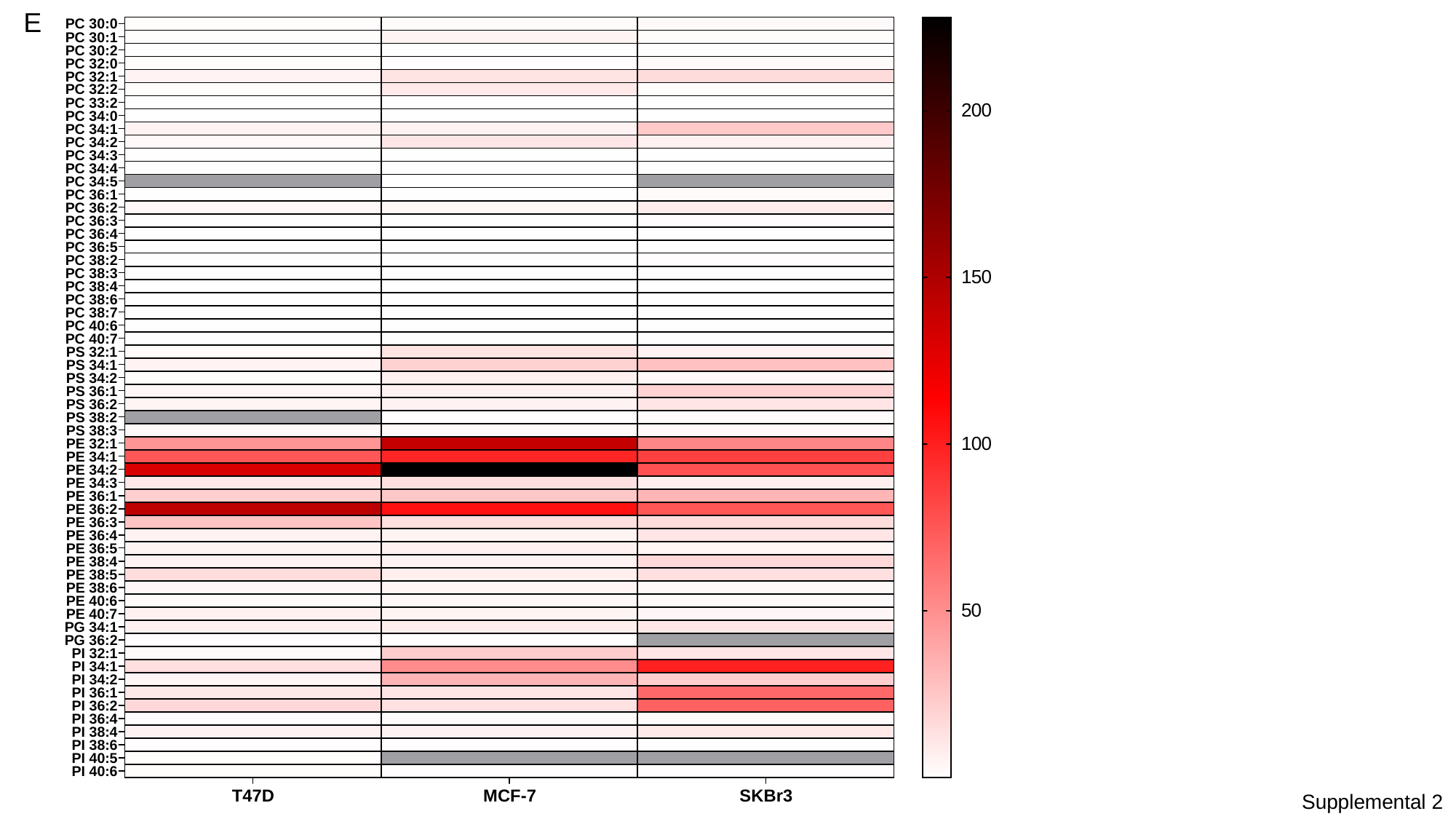

E
Supplemental 2

### Slide 6
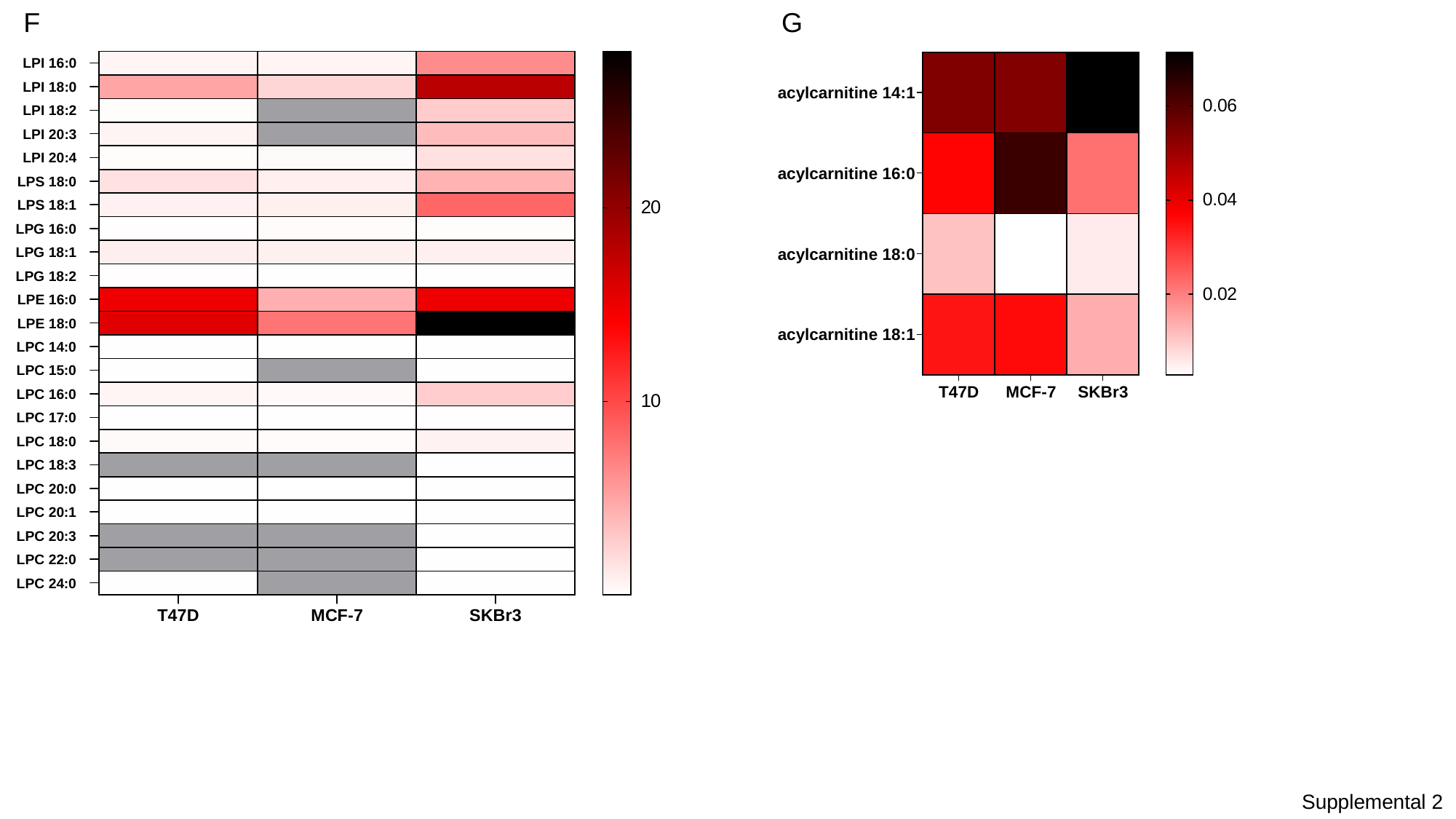

F
G
Supplemental 2

### Slide 7
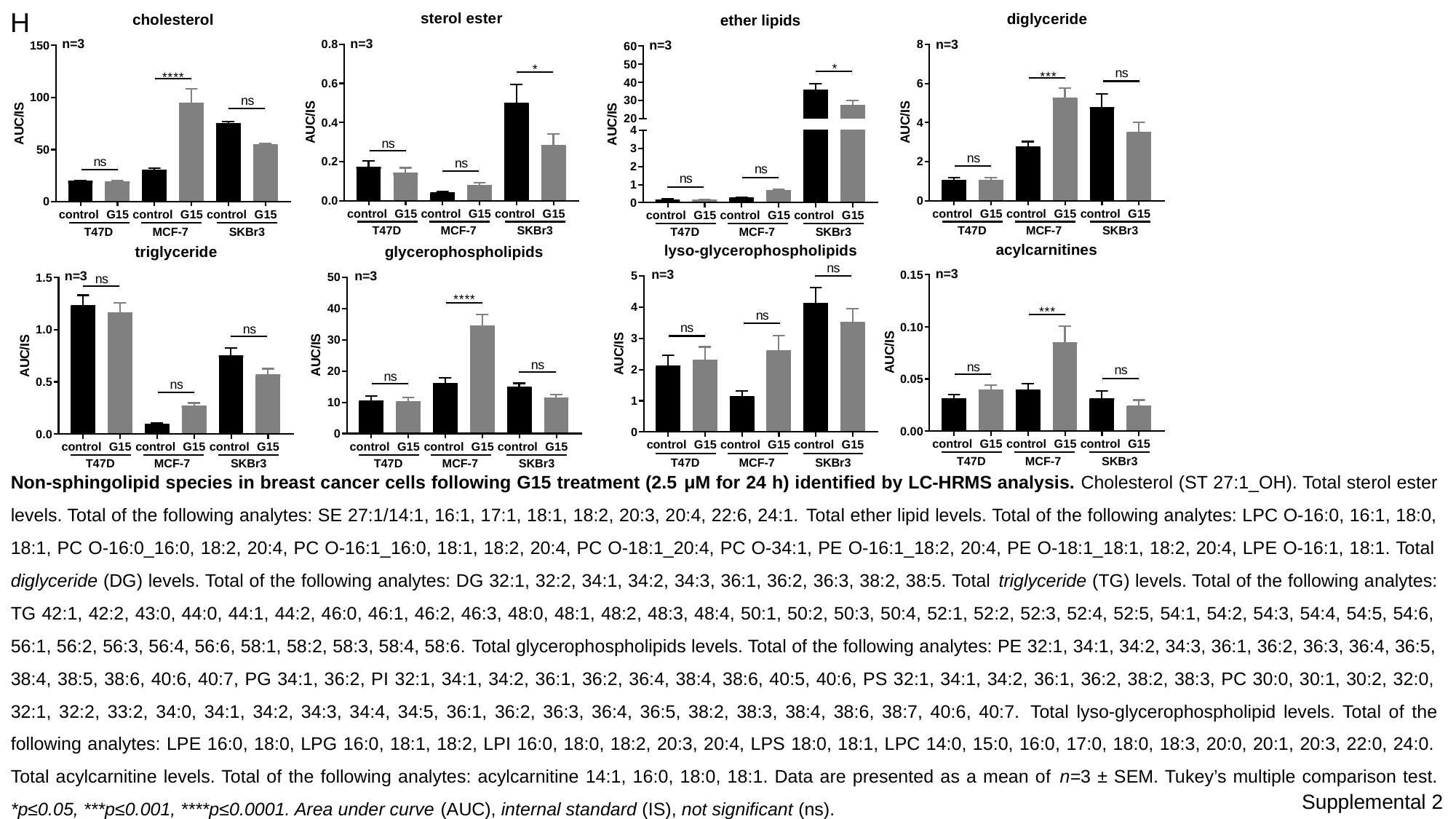

H
Non-sphingolipid species in breast cancer cells following G15 treatment (2.5 μM for 24 h) identified by LC-HRMS analysis. Cholesterol (ST 27:1_OH). Total sterol ester levels. Total of the following analytes: SE 27:1/14:1, 16:1, 17:1, 18:1, 18:2, 20:3, 20:4, 22:6, 24:1. Total ether lipid levels. Total of the following analytes: LPC O-16:0, 16:1, 18:0, 18:1, PC O-16:0_16:0, 18:2, 20:4, PC O-16:1_16:0, 18:1, 18:2, 20:4, PC O-18:1_20:4, PC O-34:1, PE O-16:1_18:2, 20:4, PE O-18:1_18:1, 18:2, 20:4, LPE O-16:1, 18:1. Total diglyceride (DG) levels. Total of the following analytes: DG 32:1, 32:2, 34:1, 34:2, 34:3, 36:1, 36:2, 36:3, 38:2, 38:5. Total triglyceride (TG) levels. Total of the following analytes: TG 42:1, 42:2, 43:0, 44:0, 44:1, 44:2, 46:0, 46:1, 46:2, 46:3, 48:0, 48:1, 48:2, 48:3, 48:4, 50:1, 50:2, 50:3, 50:4, 52:1, 52:2, 52:3, 52:4, 52:5, 54:1, 54:2, 54:3, 54:4, 54:5, 54:6, 56:1, 56:2, 56:3, 56:4, 56:6, 58:1, 58:2, 58:3, 58:4, 58:6. Total glycerophospholipids levels. Total of the following analytes: PE 32:1, 34:1, 34:2, 34:3, 36:1, 36:2, 36:3, 36:4, 36:5, 38:4, 38:5, 38:6, 40:6, 40:7, PG 34:1, 36:2, PI 32:1, 34:1, 34:2, 36:1, 36:2, 36:4, 38:4, 38:6, 40:5, 40:6, PS 32:1, 34:1, 34:2, 36:1, 36:2, 38:2, 38:3, PC 30:0, 30:1, 30:2, 32:0, 32:1, 32:2, 33:2, 34:0, 34:1, 34:2, 34:3, 34:4, 34:5, 36:1, 36:2, 36:3, 36:4, 36:5, 38:2, 38:3, 38:4, 38:6, 38:7, 40:6, 40:7. Total lyso-glycerophospholipid levels. Total of the following analytes: LPE 16:0, 18:0, LPG 16:0, 18:1, 18:2, LPI 16:0, 18:0, 18:2, 20:3, 20:4, LPS 18:0, 18:1, LPC 14:0, 15:0, 16:0, 17:0, 18:0, 18:3, 20:0, 20:1, 20:3, 22:0, 24:0. Total acylcarnitine levels. Total of the following analytes: acylcarnitine 14:1, 16:0, 18:0, 18:1. Data are presented as a mean of n=3 ± SEM. Tukey’s multiple comparison test. *p≤0.05, ***p≤0.001, ****p≤0.0001. Area under curve (AUC), internal standard (IS), not significant (ns).
Supplemental 2

### Slide 8
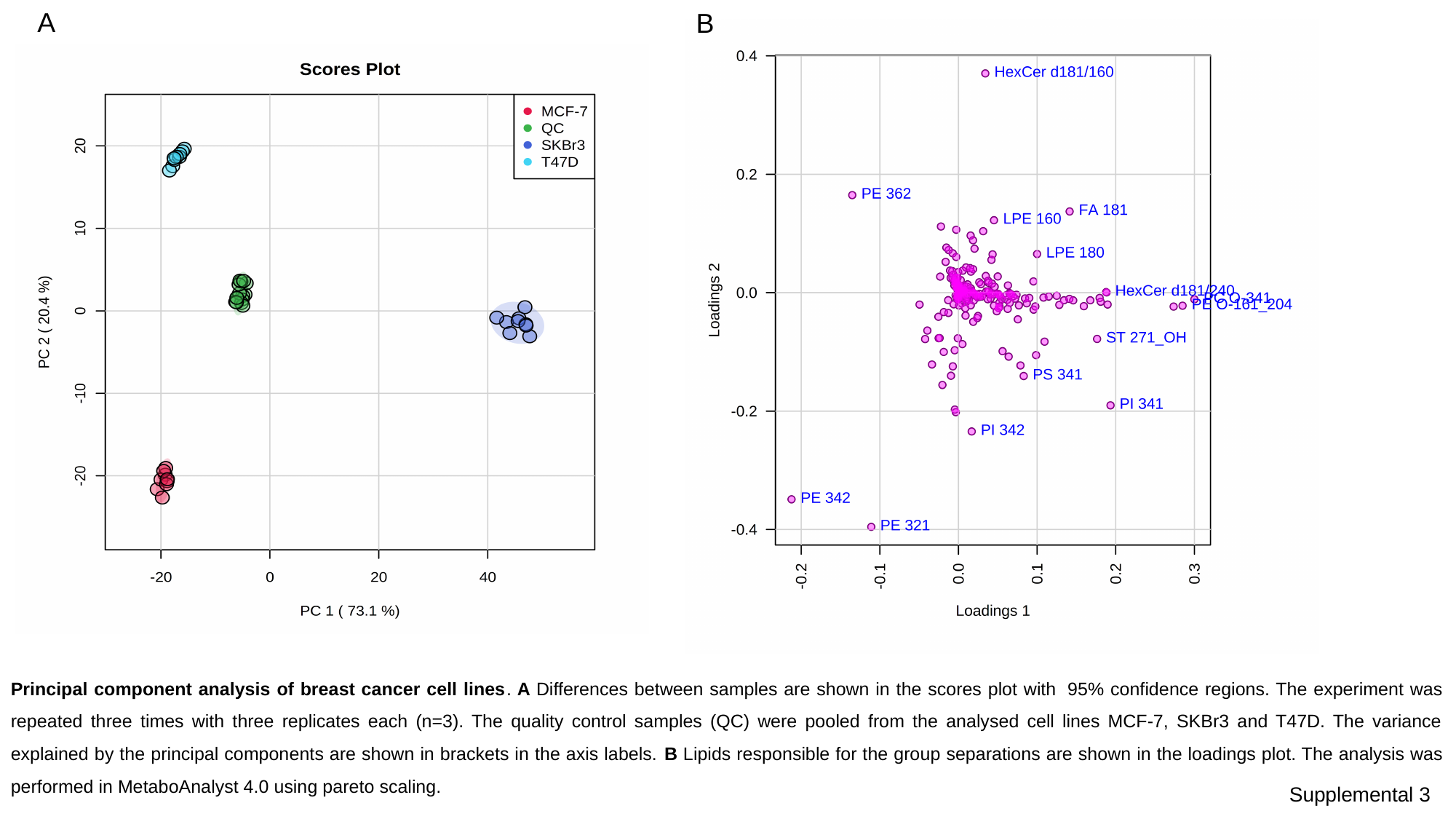

A
B
Principal component analysis of breast cancer cell lines. A Differences between samples are shown in the scores plot with  95% confidence regions. The experiment was repeated three times with three replicates each (n=3). The quality control samples (QC) were pooled from the analysed cell lines MCF-7, SKBr3 and T47D. The variance explained by the principal components are shown in brackets in the axis labels. B Lipids responsible for the group separations are shown in the loadings plot. The analysis was performed in MetaboAnalyst 4.0 using pareto scaling.
Supplemental 3
